# Oseltamivir (Tamiflu), a Commonly Prescribed Antiviral Drug, Mitigates Hearing Loss in Mice

**DOI:** 10.1101/2024.05.06.592815

**Authors:** Emma Sailor-Longsworth, Richard D. Lutze, Matthew A. Ingersoll, Regina G. Kelmann, Kristina Ly, Duane Currier, Taosheng Chen, Jian Zuo, Tal Teitz

**Affiliations:** Department of Pharmacology and Neuroscience, School of Medicine, Creighton University, Omaha, NE 68178, USA; Department of Chemical Biology and Therapeutics, St. Jude Children’s Research Hospital, Memphis, TN, 38105, USA; Department of Biomedical Sciences, School of Medicine, Creighton University, Omaha, NE 68178, USA

## Abstract

Hearing loss affects up to 10% of all people worldwide, but currently there is only one FDA-approved drug for its prevention in a subgroup of cisplatin-treated pediatric patients. Here, we performed an unbiased screen of 1,300 FDA-approved drugs for protection against cisplatin-induced cell death in an inner ear cell line, and identified oseltamivir phosphate (brand name Tamiflu), a common influenza antiviral drug, as a top candidate. Oseltamivir phosphate was found to be otoprotective by oral delivery in multiple established cisplatin and noise exposure mouse models. The drug conferred permanent hearing protection of 15-25 dB SPL for both female and male mice. Oseltamivir treatment reduced in mice outer hair cells death after cisplatin treatment and mitigated cochlear synaptopathy after noise exposure. A potential binding protein, ERK1/2, associated with inflammation, was shown to be activated with cisplatin treatment and reduced by oseltamivir cotreatment in cochlear explants. Importantly, the number of infiltrating immune cells to the cochleae in mice post noise exposure, were significantly reduced with oseltamivir treatment, suggesting an anti-inflammatory mechanism of action. Our results support oseltamivir, a widespread drug for influenza with low side effects, as a promising otoprotective therapeutic candidate in both cisplatin chemotherapy and traumatic noise exposure.

## Introduction

Hearing loss is a highly prevalent cause of global disability and is estimated to affect up to 10% of the population worldwide (Carroll et al., 2017; Natarajan et al., 2023; Oishi and Schacht, 2011). Hearing dysfunction negatively impacts an individual’s quality of life and is a known risk factor for poor academic performance, dementia, overall cognitive decline and depression (Lin et al., 2023; Loughrey et al., 2018; Orgel et al., 2016). Among the most common causes of severe hearing loss are cisplatin, a broadly used chemotherapeutic agent, and high-decibel noise exposure (Callejo et al., 2015; Natarajan et al., 2023). Up to 60% of patients treated with cisplatin develop severe, permanent hearing loss and noise-induced hearing loss is the second most frequent form of sensorineural hearing impairment after age-related hearing loss (Callejo et al., 2015; Carroll et al., 2017; Dobie, 2008). In 2022, sodium thiosulfate (STS) (brand name PEDMARK®) received FDA approval for the prevention of cisplatin-induced hearing loss in pediatric patients with localized, nonmetastatic solid tumors (Brock et al., 2023; Dhillon, 2023; Freyer et al., 2023). To date, it is the only FDA-approved drug available to prevent hearing loss by any cause (Dhillon, 2023). STS has been a great advancement in the field for protection from hearing loss in pediatric patients; however, like all drugs, it has limitations to the patient population that can be treated with the drug (Brock et al., 2023; Hazlitt et al., 2018; Orgel et al., 2022). Consequently, there is a dire need to find more drugs that protect from noise-and cisplatin-induced hearing loss for metastatic pediatric cancer patients and the adult cancer population.

Our laboratory and others have demonstrated the existence of several molecular pathways common to cisplatin ototoxicity and acoustic trauma (Hazlitt et al., 2018; Ingersoll et al., 2024, 2023, 2020; Lutze et al., 2023; Teitz et al., 2018). This suggests that targeting shared molecular pathways could protect from both insults of hearing loss. For example, we have shown that mitogen activated protein kinase (MAPK) pathway activation occurs with both cisplatin and damaging noise exposure in the supporting and hair cells of the inner ear, and multiple inhibitors of the MAPK pathway shield mice from both cisplatin- and noise-induced hearing loss (Ingersoll et al., 2024, 2023, 2020; Lutze et al., 2023). In addition, molecular processes such as activation of shared programmed cell death pathways and enhanced inflammation with immune cell infiltration occur following both insults (Frye et al., 2019; He et al., 2020; Ramkumar et al., 2021; Wood and Zuo, 2017; Wu et al., 2020). Depending on the intensity of the noise exposure and levels of cisplatin damage, some of the same inner ear cell types and neuron fibers that are damaged by cisplatin are damaged by noise exposure, including the outer hair cells (OHCs), supporting cells, the stria vascularis, auditory nerve synapses, and the spiral ganglion neurons (Kros and Steyger, 2019; Kurabi et al., 2017; Sheth et al., 2017; Wang et al., 2023). Therapeutics that target these overlapping molecular and cellular events may therefore protect against both insults in the inner ear.

We first identified oseltamivir phosphate (brand name Tamiflu®) in a high-throughput screen of 1,300 FDA approved drugs for protection against cisplatin-induced cell death performed with an inner ear cell line HEI-OC1 (Ingersoll et al., 2020; Teitz et al., 2018, 2016). Oseltamivir is uniquely well-positioned for drug repurposing for hearing protection. In general, repurposing FDA-approved drugs has advantages over the development of new chemical entities due to significant reductions in time and cost to achieve regulatory approval (Kumar et al., 2019). Oseltamivir, specifically, is an FDA-approved oral drug since 1999 for the treatment of acute, uncomplicated influenza in patients 2 weeks of age and older whose flu symptoms have not lasted more than two days (Dutkowski et al., 2003; He et al., 1999; McClellan and Perry, 2001). The drug is approved to treat type A and B influenza and for the prevention of influenza in adults and children aged one year and older (Dutkowski et al., 2003; McClellan and Perry, 2001). Oseltamivir inhibits viral neuraminidases required to cleave budding virions from the host cell, blocking further viral replication (Moscona, 2005). The drug has low toxicity and high tolerability in the general population with nausea and vomiting as possible side effects in the first two days of treatment that can be reduced by taking the drug with food (McClellan and Perry, 2001). Extremely rare side effects of sudden confusion, delirium, hallucinations, unusual behavior, or self-injury have been reported in children and young adults from Asian populations although the association with the drug is not clear (Chen et al., 2019; Ho et al., 2010; Izumi et al., 2007; Morimoto et al., 2015). The prodrug oseltamivir phosphate has the advantage of being highly orally bioavailable, at least 75% of the orally administered drug reaches the systemic circulation in humans and mice in the form of the active metabolite oseltamivir carboxylate (Davies, 2010; Dutkowski et al., 2003; He et al., 1999). The half-life of oseltamivir phosphate after oral administration is 2 to 3 hours in mice and humans, and the half-life of oseltamivir carboxylate is 2 to 3 hours in mice and 6 to 10 hours in humans (He et al., 1999; Honce et al., 2023.; Pillai et al., 2015; Shin et al., 2017).

In this study, we demonstrate oseltamivir phosphate’s ability to protect from cisplatin and noise-induced hearing loss by oral delivery in various preclinical mouse models. Both the prodrug, oseltamivir phosphate, and its antiviral active form, oseltamivir carboxylate, were tested for protection from cisplatin-induced outer hair cell death in cochlear explants and the drug interference with cisplatin’s tumor killing ability was measured in tumor cell lines. We tested the drug’s ability to protect from both cisplatin and noise-induced hearing loss in multiple mouse models and optimized the treatment protocol for oseltamivir treatment. Next, we examined the top mammalian molecular target of both oseltamivir phosphate and oseltamivir carboxylate, predicted by structure binding analysis, ERK1/2, and found that MAPK activation was upregulated in cisplatin-treated cochlear explants and reduced with oseltamivir treatment. Finally, immune cell infiltration was measured following noise exposure and oseltamivir treatment to determine whether the drug reduces inflammation following noise insult and elucidate the potential mechanisms of protection of oseltamivir. Overall, the results of this study demonstrate that oseltamivir, a widely used antiviral drug, is a promising therapeutic candidate for protection from sensorineural hearing impairment caused by both cisplatin and noise insults.

## Results

### Oseltamivir phosphate and its active form, oseltamivir carboxylate, protect against cisplatin-induced hair cell loss in mouse cochlear explants

In the aim of identifying efficient and safe drugs for protection from cisplatin and noise-induced hearing loss, we conducted unbiased high-throughput screens of 1,300 FDA-approved drugs for cisplatin-induced cell death protection in an inner ear cell line HEI-OC1, employing the methodology previously described (Ingersoll et al., 2020; Teitz et al., 2018, 2016). The prodrug, oseltamivir phosphate, tested at dose of 3 µM, was a top hit in the screen reducing 95% of the caspase-3/7 cell death activity of the cisplatin-treated cells. To confirm the drug’s ability to protect from OHCs death, we tested the prodrug, oseltamivir phosphate, and the active antiviral drug, oseltamivir carboxylate, in FVB/NJ mouse P3 cochlear explants. Briefly, P3 pups’ cochleae were dissected and cultured in media. Explants were then treated with medium alone, oseltamivir phosphate or carboxylate alone, 150 μM cisplatin alone, or varying of doses of oseltamivir phosphate or carboxylate with 150 μM cisplatin. 24 hours after treatment, samples were fixed and stained with phalloidin to count the number of hair cells per 160 μm for each sample. Oseltamivir phosphate protected from cisplatin induced outer hair cell death with an EC_50_ of 450 nM and 1 μM oseltamivir phosphate combined with cisplatin had a similar number of hair cells compared to medium alone (Figure 1A & 1D). In addition, none of the oseltamivir phosphate doses tested were toxic to explants, and the therapeutic index was greater than 220. For the active form of oseltamivir, oseltamivir carboxylate (Figure 1B), we found that the EC_50_ of protection from outer hair cell death was 505 nM which is very close to the prodrug’s EC_50_ (Figure 1C & E). The LD_50_ and therapeutic index of the active form of oseltamivir were not changed as well, compared to the prodrug (Figure 1). This indicated that both oseltamivir’s prodrug and its active metabolite have similar efficacy in the context of protection from OHCs death in the cochlear explants.

**Figure 1:**
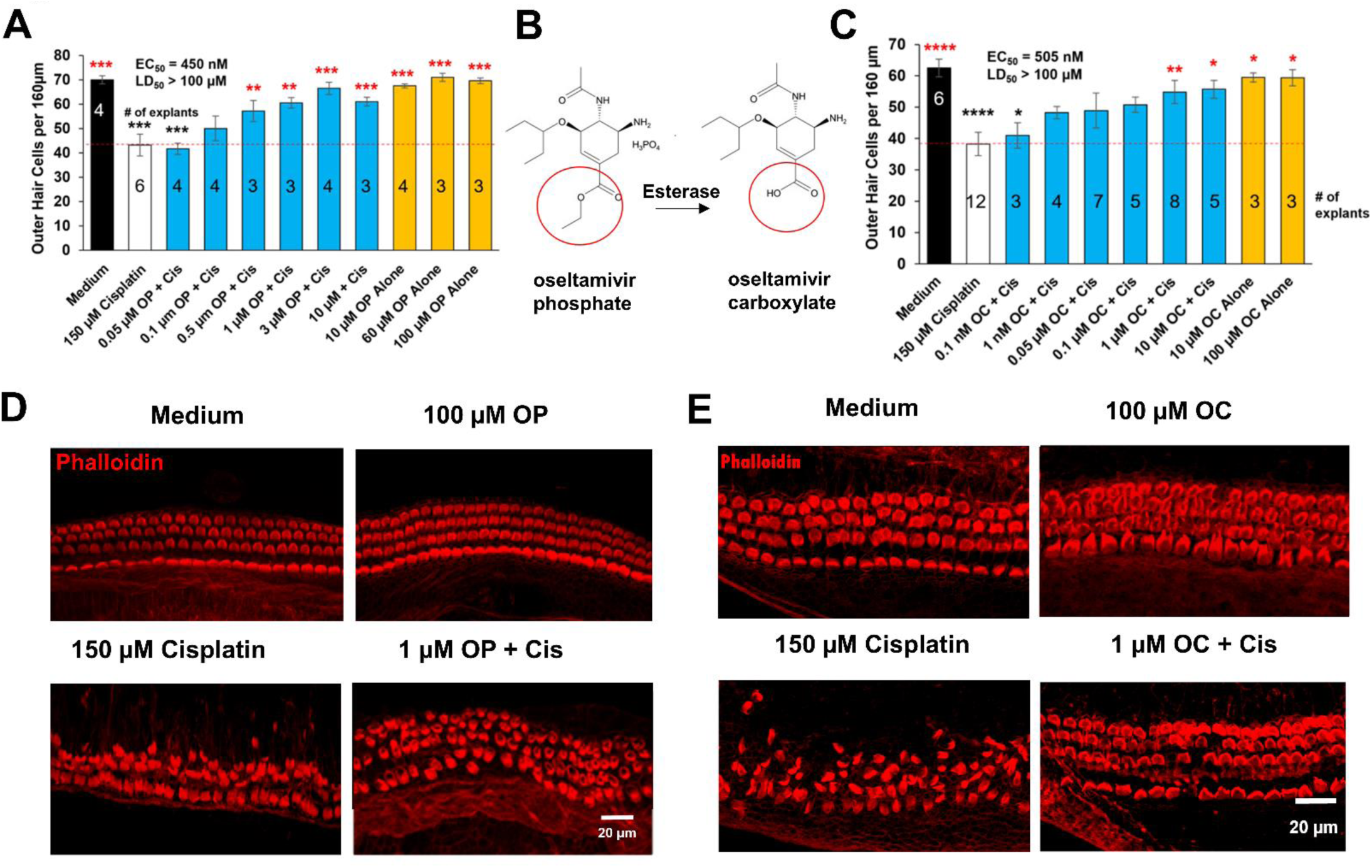
Oseltamivir, an antiviral neuraminidase inhibitor, reduces cisplatin-induced cell death in murine cochlear explants. **(A)** Dose-response relationship of the pro-drug, oseltamivir phosphate, and outer hair cell count in P3 FVB pup cochlear explants treated with cisplatin. Black: media alone; white: cisplatin alone, gold: oseltamivir phosphate alone; blue: cisplatin (150 μM) and oseltamivir phosphate (0.05—10 μM) cotreatment. Cotreated samples were pretreated with oseltamivir phosphate for one hour prior to cisplatin treatment, and then cultured in oseltamivir phosphate- and cisplatin-treated media for 24 hours. Numbers within bars denote # of explants per treatment. Samples were then stained with phalloidin with outer hair cell (OHC) counts per 160 μm taken from the cochlear middle turn. **(B)** Molecular structure of the prodrug, oseltamivir phosphate, and its active antiviral metabolite, oseltamivir carboxylate. Hydrolysis of the circled ether group by hepatic esterase yields oseltamivir carboxylate, a sialic acid analogue and viral neuraminidase inhibitor. **(C)** Dose-response relationship of the metabolite oseltamivir carboxylate and outer hair cell count in P3 FVB pup cochlear explants treated with cisplatin. Black: media alone; white: cisplatin alone, gold: oseltamivir carboxylate alone; blue: cisplatin (150 μM) and oseltamivir carboxylate (0.001—10 μM) cotreatment. **(D-E)** Representative confocal images of explants treated with medium alone, 100 μM oseltamivir phosphate (D) or oseltamivir carboxylate (E), 150 μM cisplatin, and 3 μM oseltamivir phosphate (d) or 1 μM oseltamivir carboxylate (E) + 150 μM cisplatin. Data shown as means ± SEM*, *p* < 0.05, \*\**p* < 0.01, \*\*\**p* < 0.001 compared to cisplatin alone (red) and medium alone (black) by one-way analysis of variance (ANOVA) with Bonferroni post hoc test.

### Oseltamivir does not interfere with cisplatin’s tumor killing efficacy in lung carcinoma and neuroblastoma cell lines

Six different cell lines were treated with cisplatin and oseltamivir to determine whether the drug interferes with cisplatin’s ability to kill cancer cells. Three non-small cell lung carcinoma (H1155, SHP-77, and A549) and three neuroblastoma cell lines (Kelly, SK-N-AS, and SH-SY5Y) were utilized as these tumors are commonly treated with cisplatin (Ghosh, 2019). Each cell line was plated into 96 well plates and every treatment group were in replicates of six. Each cell line was treated with a concentration of cisplatin that killed approximately 30-50% of cells and six different doses of oseltamivir were tested. A dose of 30 μM oseltamivir phosphate with five serial dilutions of 1:3 were tested to ensure a wide range of doses does not interfere with cisplatin. Every dose of oseltamivir in all six cell lines did not interfere with cisplatin’s killing efficacy as determined by the Cell Titer-Glo assay (Figure 2). Every cell line treated with oseltamivir and cisplatin did not have significantly different cell viability compared to the cisplatin alone treated wells with any of the oseltamivir doses tested.

**Figure 2:**
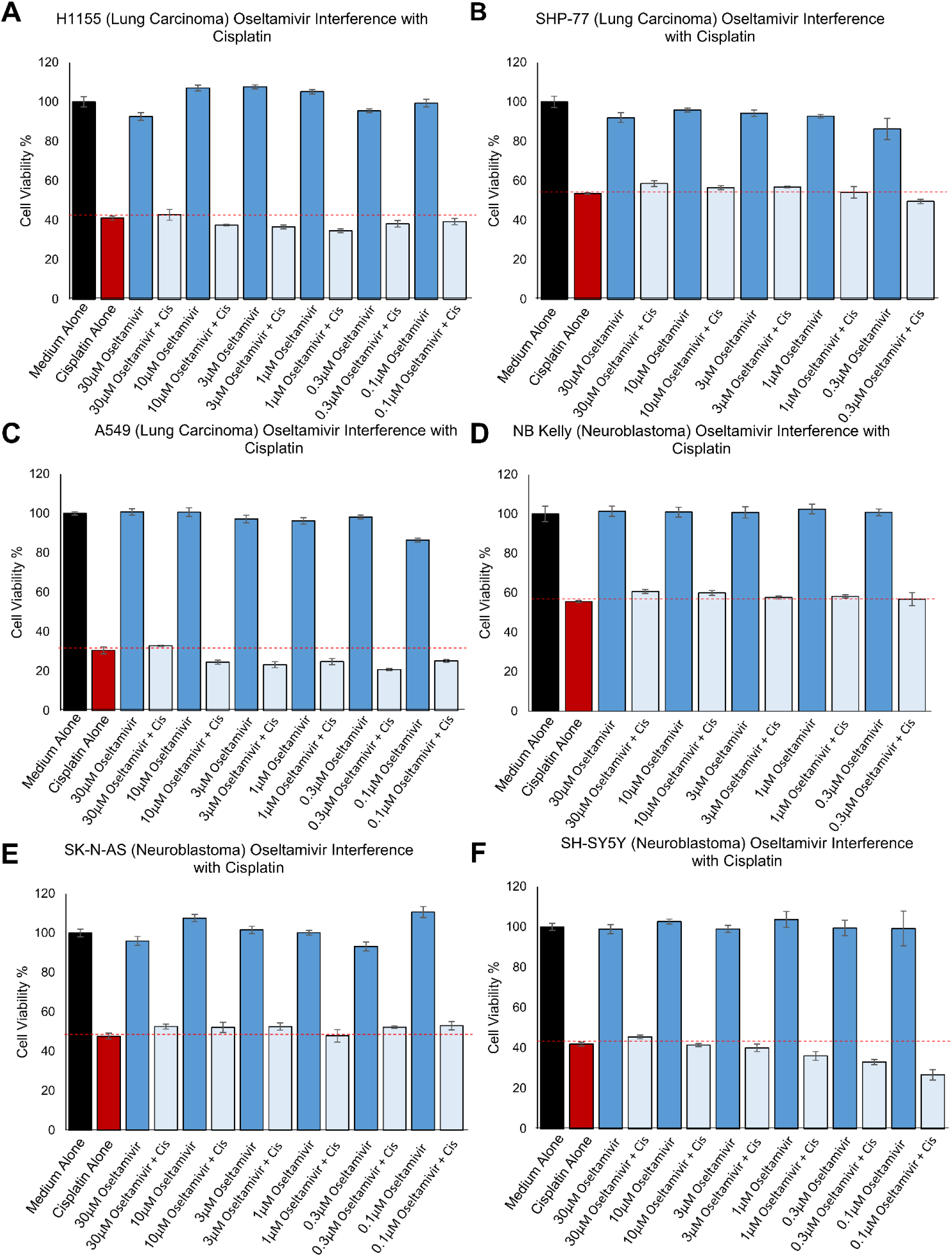
Oseltamivir does not interfere with cisplatin’s tumor killing ability in 3 lung carcinoma and 3 neuroblastoma cell lines. 3 neuroblastoma and 3 lung carcinoma cell lines were treated with cisplatin and 6 varying oseltamivir concentrations and then the Cell Titer-Glo assay was performed to determine cell viability. Cell Viability graphs after cisplatin and oseltamivir treatment for the **(A)** H1155 **(B)** SHP-77 **(C)** A549 **(D)** Kelly **(E)** SK-N-AS and **(F)** SH-SY5Y cell lines are shown. Medium Alone (Black), Cisplatin Alone (Red), oseltamivir Alone (Dark Blue), and oseltamivir + Cisplatin (Light Blue). Data shown as means ± SEM, compared to cisplatin alone by one-way ANOVA with Bonferroni post-hoc test. n=6 wells

### Oseltamivir protects FVB/NJ mice from cisplatin ototoxicity after a single, high dose of cisplatin

To test whether oseltamivir protects from cisplatin ototoxicity *in* vivo, FVB/NJ mice were treated orally with 50 mg/kg oseltamivir phosphate, 45 minutes before the administration of 30 mg/kg cisplatin. Then, mice were treated with oseltamivir later that day and treatment continued for a total of three days, twice a day (Figure 3A). Mice cotreated with oseltamivir and cisplatin had significantly lower ABR threshold shifts at the 32 kHz region compared to mice only treated with cisplatin (Figure 3B & C). Oseltamivir cotreated mice had an ABR threshold shift reduction of 15 dB compared to the cisplatin alone treatment. Mouse cochleae were then collected and stained with myosin VI to count the number of OHCs per 160 µm (Figure 3D). Cotreatment with oseltamivir significantly reduced OHCs death at the middle and basal regions compared to cisplatin alone treatment (Figure 3D & E). Cisplatin alone treated mice had an average of 52 OHCs at middle region and 43 at the basal region. Oseltamivir and cisplatin cotreated mice had an average 64 and 59 OHCs at the middle and basal regions, respectively.

**Figure 3:**
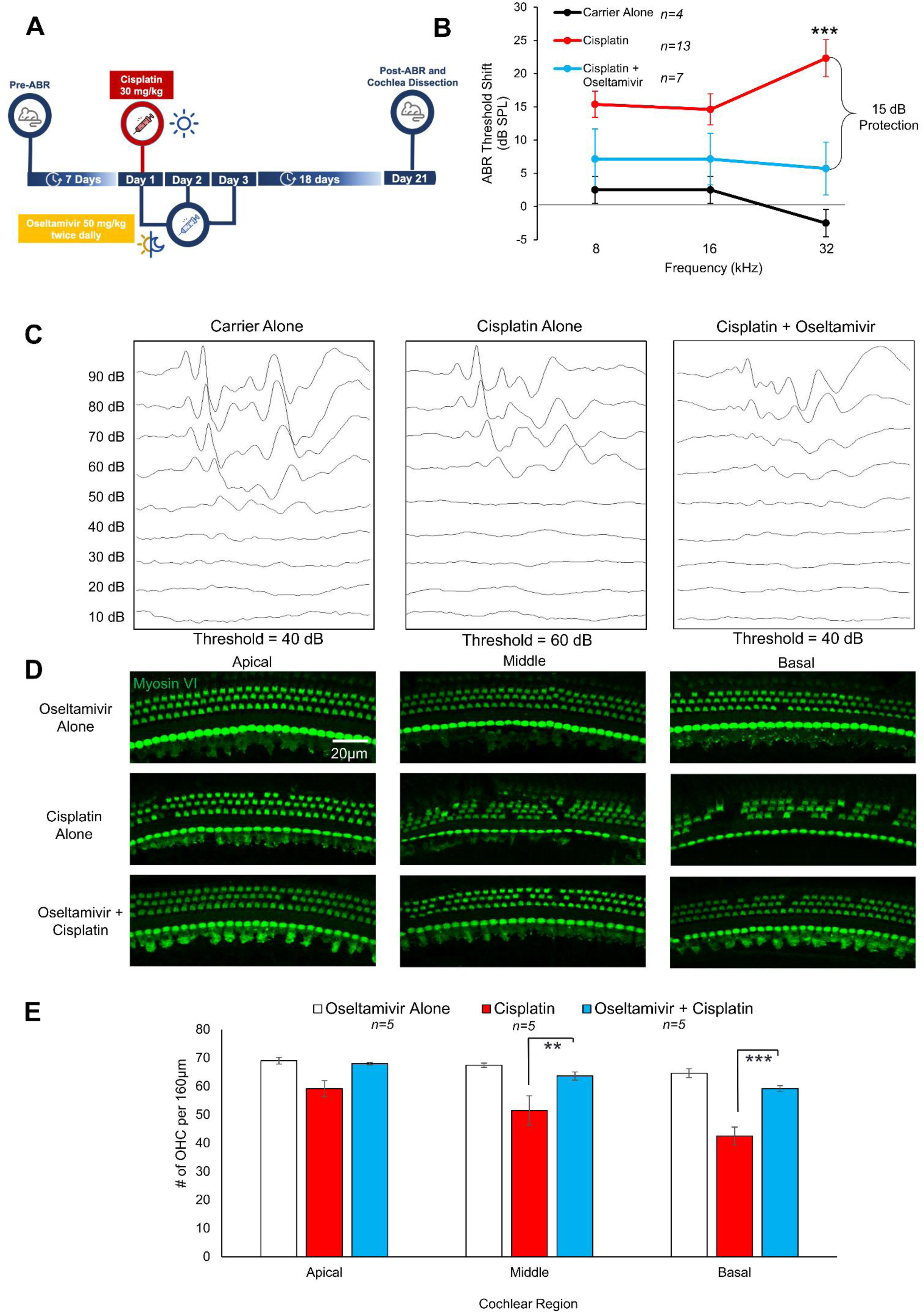
Oseltamivir protects mice from cisplatin-induced hearing loss and OHC loss after a single, high dose of cisplatin. **(A)** Treatment schedule in which mice were treated with a single dose of 30 mg/kg cisplatin and 50 mg/kg oseltamivir twice a day, once in morning and once at night, for three days. **(B)** ABR threshold shifts following treatment protocol in (A). Carrier Alone (black), Cisplatin Alone (Red), and oseltamivir + Cisplatin (Blue). Data shown as means ± SEM, compared to cisplatin alone by two-way ANOVA with Bonferroni post hoc test. ***p<0.001 **(C)** Representative ABR traces after the treatment protocol in (A) at the 32 kHz frequency of carrier alone, cisplatin alone, and cisplatin + oseltamivir treated mice. **(D)** Representative whole mount cochlear images stained with myosin VI following treatment protocol in (A). Cochlear whole mount sections from the apical, middle, and basal regions were imaged. **(E)** Quantification of images shown in (C). The number of OHCs per 160 μm was counted per sample in each region. Oseltamivir Alone (White), Cisplatin Alone (Red), and oseltamivir + Cisplatin (Blue). Data shown as means ± SEM, compared to cisplatin alone by one-way ANOVA with Bonferroni post hoc test., **p<0.01, ***p<0.001.

### Oseltamivir protects mice from cisplatin-induced hearing loss in a clinically relevant, multicycle cisplatin protocol with low dose of 10 mg/kg/bw

Patients are not given a single high dose of cisplatin; therefore, we utilized a clinically relevant mouse model that mimics cisplatin treatment in humans (Fernandez et al., 2019; Ingersoll et al., 2023; Roy et al., 2013). Figure 4A shows an outline of this protocol in which mice were treated orally with 50, 10, or 2 mg/kg oseltamivir 45 minutes before the cisplatin treatment in the morning. 3 mg/kg cisplatin was then administered via intraperitoneal (IP) injection and another dose of oseltamivir was given later that day. Cisplatin treatment occurred for a total of four consecutive days and oseltamivir was administered for a total of five consecutive days. After the treatment period, no treatments occurred for nine days and then this treatment cycle was repeated for a total of three times. Mice treated with 50 and 10 mg/kg oseltamivir had significantly lower ABR threshold shifts at the 16 and 32 kHz regions compared to cisplatin alone treated mice (Figure 4B & C). Mice cotreated with oseltamivir and cisplatin had an average ABR threshold shift reduction of 15 dB at the 16 and 32 kHz regions compared to cisplatin alone treatment. 2 mg/kg oseltamivir treatment had the same ABR threshold shift as the cisplatin alone treatment (Figure 4B). 50 and 10 mg/kg oseltamivir cotreated mice with cisplatin had significantly higher ABR wave 1 amplitudes at 90 dB compared to cisplatin alone treated mice (Figure 4D) but no reduction in DPOAE threshold shifts was observed with any dose of oseltamivir treatment (Figure 4E). The 50 and 10 mg/kg oseltamivir treatment conferred a significant reduction in OHCs loss at the apical and middle regions compared to cisplatin alone (Figure 4F & G). Cisplatin alone treated mice had an average of 32 OHCs per 160μm at the apical region and 9 OHCs at the middle region while mice cotreated with 50 mg/kg oseltamivir and cisplatin had 53 and 33 at the apical and middle regions, respectively (Figure 4F & G). Mice cotreated with 10 mg/kg oseltamivir and cisplatin had an average of 59 OHCs at the apical region and 37 OHCs at the middle region (Figure 4F & G). There was no significant reduction in OHCs loss at the basal region for either oseltamivir treatment (Figure 4F & G). There was no significant difference in weight loss between the cisplatin alone treated cohort and either one of the oseltamivir and cisplatin treated groups (Figure 4H).

**Figure 4:**
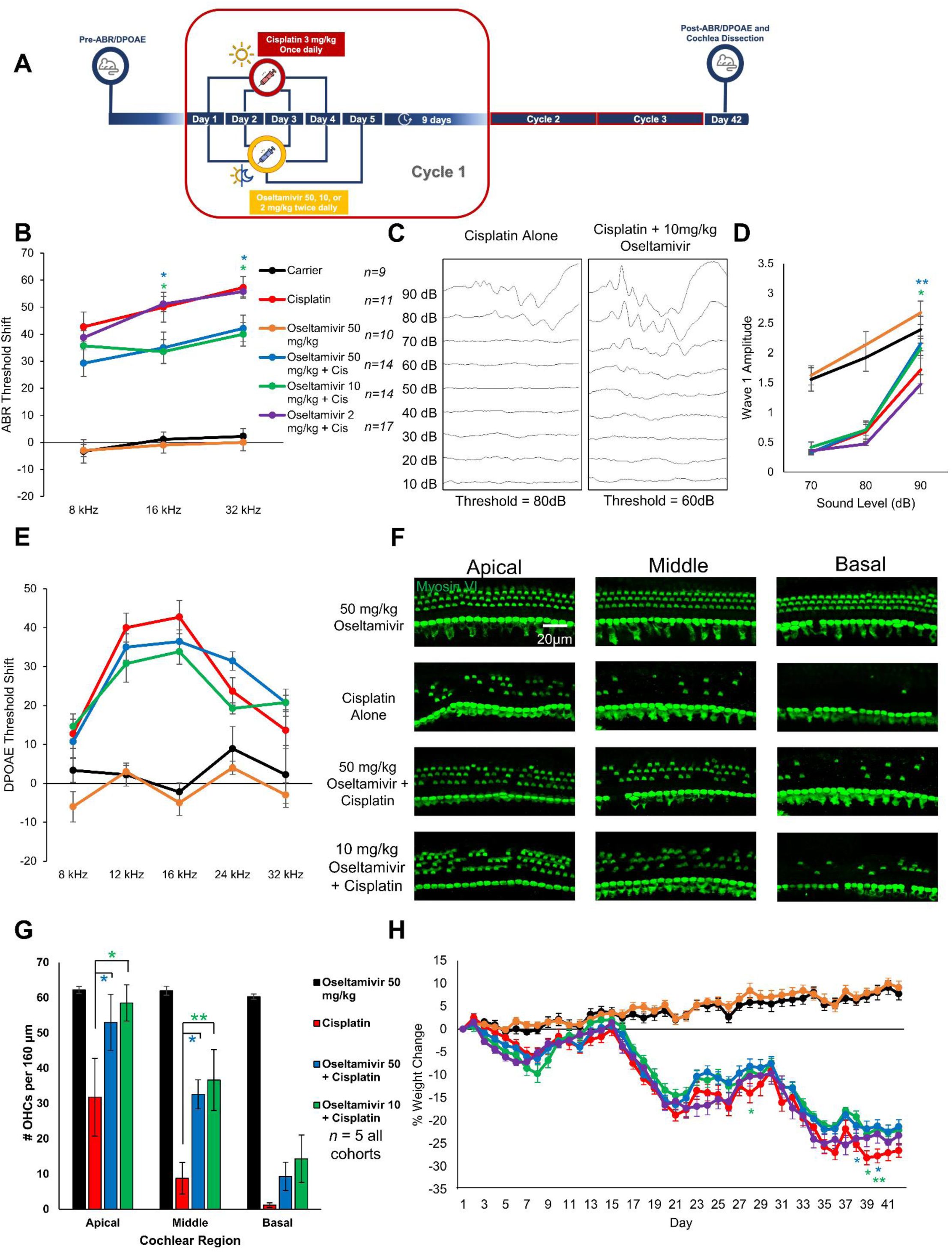
Oseltamivir protects mice from cisplatin-induced hearing loss and OHC death in a clinically relevant, multicycle cisplatin treatment protocol. **(A)** Treatment protocol where mice were treated with 3 mg/kg in the morning for 4 days and then treated with 50, 10, or 2 mg/kg oseltamivir in the morning and night for 5 days. A 9-day recovery period occurred and then this cycle was repeated two more times for a total of 3 cycles. **(B)** ABR threshold shifts following protocol in (A). **(C)** Representative ABR traces after the treatment protocol in (A) at the 32 kHz frequency of cisplatin alone and cisplatin + 10 mg/kg oseltamivir treated mice. **(D)** ABR wave one amplitude from ABR’s in (B) at the 16 kHz frequency. **(E)** DPOAE threshold shifts following treatment protocol in (A). Carrier (Black), oseltamivir Alone (Orange), Cisplatin Alone (Red), 50 mg/kg oseltamivir + Cisplatin (Blue), 10 mg/kg oseltamivir + Cisplatin (Green), and 2 mg/kg oseltamivir + Cisplatin (Purple). Data shown as means ± SEM, compared to cisplatin alone by two-way ANOVA with Bonferroni post hoc test. n=9-17. **(F)** Representative whole mount cochlear images stained with myosin VI following treatment protocol in (A). n=5 **(G)** Quantification of OHC counts per 160 μm of images shown in (E). oseltamivir Alone (Black), Cisplatin Alone (Red), 50 mg/kg oseltamivir + Cisplatin (Blue), and 10 mg/kg oseltamivir + Cisplatin (Green). Data shown as means ± SEM, compared to cisplatin alone by one-way ANOVA with Bonferroni post-hoc test. *p<0.05, **p<0.01 **(H)** Percent weight loss of all treatment groups following treatment protocol in (A). Mice were weighed everyday throughout the 42-day treatment protocol. Data shown as means ± SEM, compared to cisplatin alone by one-way ANOVA with Bonferroni post-hoc test. *p<0.05, **p<0.01, n=9-17 mice.

### Oral oseltamivir therapy protects against noise-induced ABR threshold shifts at doses of 100 and 50 mg/kg

Oseltamivir’s efficacy against noise-induced hearing loss was tested in a mouse model of acoustic trauma (Figure 5A). Briefly, ABR measurements were performed in 6-8 weeks old FVBN/J mice to ascertain baseline hearing capability. One week after baseline ABR measurements were performed, mice were subjected to a noise challenge consisting of exposure to 100 dB noise over an 8-16 kHz octave band for two hours. Beginning 24 hours after exposure, mice were treated twice daily via oral gavage with oseltamivir at 100 mg/kg, 50 mg/kg, or 10 mg/kg or with carrier alone. The treatment was terminated after three days and ABR was again measured 14 days after the noise challenge. Mice who received 100 mg/kg oseltamivir phosphate after noise exposure exhibited significantly reduced ABR threshold shifts in all tested frequencies relative to noise-exposed mice treated with carrier (average threshold shifts of 14,19, and 17 dB SPL vs 39, 36, and 35 dB SPL at 8, 16, and 32 kHz, respectively; Figure 5B & C). Both female and male mice treated with oseltamivir following noise exposure had significantly lower ABR threshold shifts compared to noise alone mice (Figure 5D). Mice that received 50 mg/kg had significant protection at 8 kHz, with an average reduction in the threshold shift which equaled 23 dB SPL (Figure 5E). No protection from noise-induced ABR threshold shifts was observed in the 10 mg/kg treatment group relative to noise-exposed controls (Figure 5F). Interestingly, no treatment group exhibited protection from DPOAE threshold shifts (Figure 5G-I).

**Figure 5:**
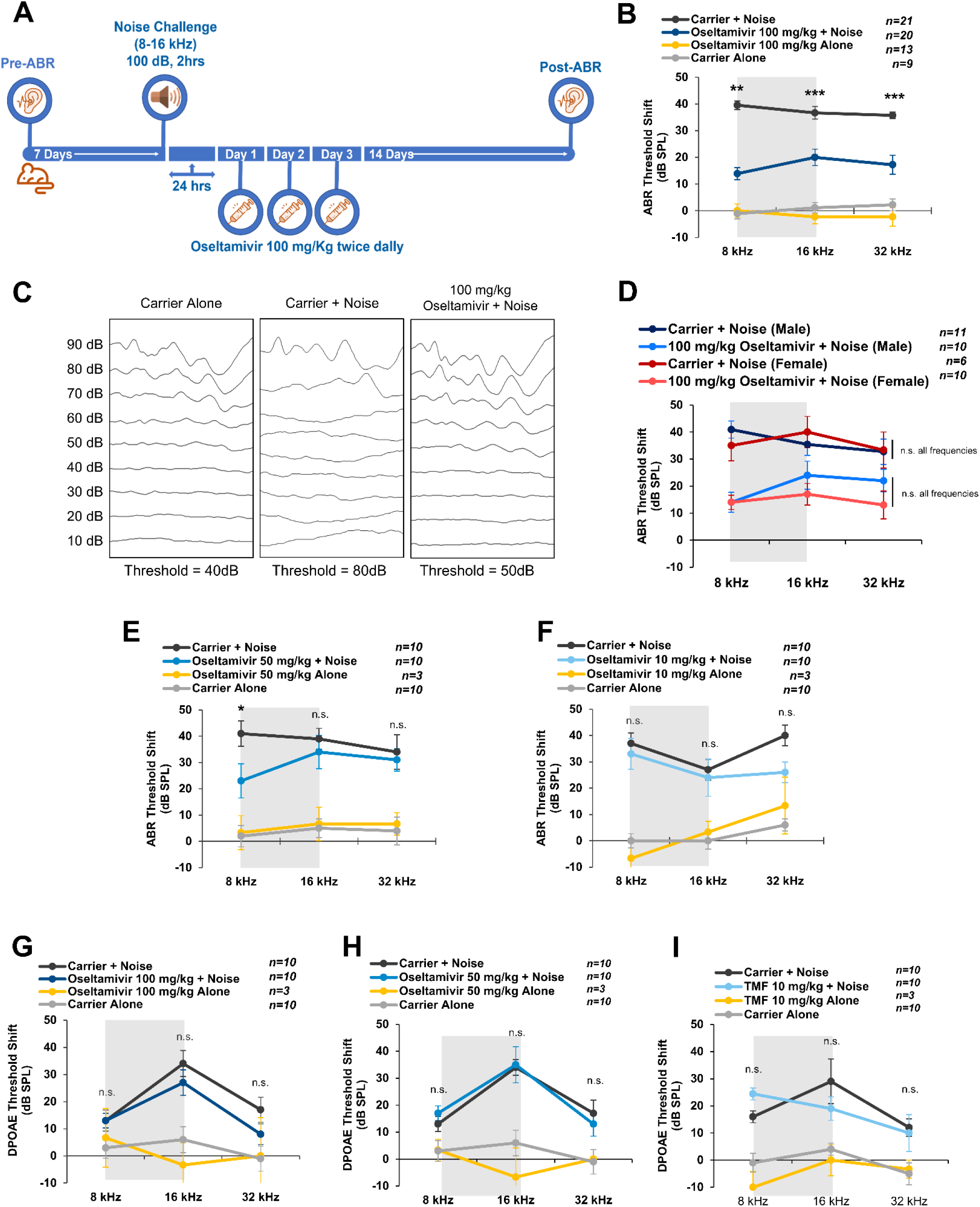
100 mg/kg and 50 mg/kg oral oseltamivir therapy protects against noise-induced ABR threshold shift, but does not rescue DPOAE. **(A)** Schematic of the treatment schedule. **(B)** 100 mg/kg oseltamivir treatment significantly protected hearing in noise-exposed mice with an average reduction of ABR threshold shift of 20-25 dB SPL at 8, 16 and 32 kHz. Data shown as means ± SEM compared to carrier + noise by two-way ANOVA with Bonferroni post-hoc test. **p<0.01, ***p<0.001. **(C)** Representative ABR traces after the treatment protocol in (A) at the 16 kHz frequency of carrier alone, carrier + noise alone, and noise + oseltamivir treated mice. **(D)** Results in (B) broken down by animal sex. **(E)** Mice treated with 50 mg/kg oseltamivir exhibited significantly lower ABR threshold shifts at 8 kHz when compared to noise-exposed mice treated with carrier alone. Data shown as means ± SEM compared to carrier + noise by two-way ANOVA with Bonferroni post-hoc test. *p<0.05. **(F)** No protection was observed with 10 mg/kg oseltamivir. **(G, H, I)** DPOAE threshold shifts were also measured for mice treated with 100 mg/kg, 50 mg/kg, and 10 mg/kg, respectively; no significant protection was observed. Data shown as means ± SEM compared to carrier + noise by two-way ANOVA with Bonferroni post-hoc test.

### Three-day oseltamivir treatment is sufficient for otoprotective effects when initiated up to 24 hours after 100 dB, 2 h noise exposure, but no protection was achieved with 106 dB, 2h noise insult

We tested whether extending treatment duration from three to five days would provide superior protection. Mice who received 100 mg/kg oseltamivir twice daily for five days beginning 24 hours’ post-noise exposure demonstrated robust protection at all frequencies (Figure 6A & B; average threshold shifts of 12-, 19-, and 26-dB SPL for oseltamivir treated mice exposed to noise vs 49-, 39- and 46 dB SPL for carrier treated noise exposed mice at 8, 16, and 32 kHz, respectively). These results were not statistically superior to mice who received three days of treatment (95% CI = −13, 23; −14, 22; and −15, 23 for 8, 16, and 32 kHz, respectively) indicating that a three-day treatment duration is sufficient to achieve protection against elevated ABR threshold shifts.

**Figure 6:**
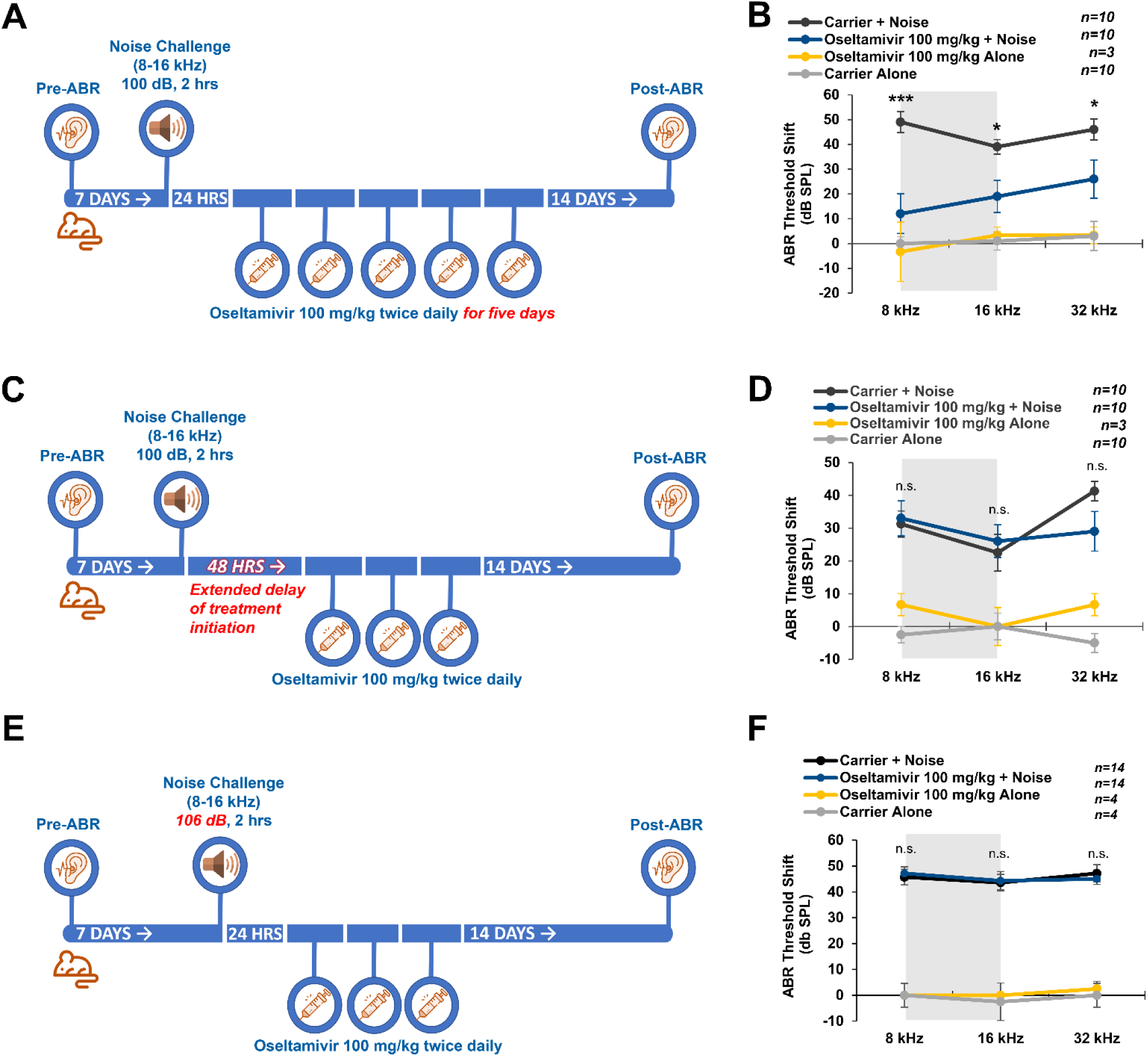
100 mg/kg oral oseltamivir treatment twice daily is protective against ABR threshold shifts when started 24 hours after noise and continued at least for three days, but not when started 48 hours post-noise challenge or following 106 dB noise exposure. **(A)** Schematic diagram of extended treatment schedule. Oseltamivir was initially tested at 100 mg/kg twice daily, beginning 24 hours post noise challenge and continuing for three days. The first alternative treatment schedule involved extending treatment to 5 days**. (B)** ABR threshold shifts for mice treated with oseltamivir for an extended duration of five days. **(C)** Schematic diagram of delayed treatment schedule (48 hrs post-noise exposure). **(D)** ABR threshold shifts for mice treatment with oseltamivir (100 mg/kg) beginning 48 hours post-noise exposure. **(E)** Schematic diagram illustrating treatment schedule following a noise exposure of increased volume (from 100 to 106 dB). Mice were treated as in Figure 5A with no adjustment made in treatment timing or duration. **(F)** ABR threshold shifts for mice treated with 100 mg/kg oseltamivir phosphate following 106 dB noise challenge. Data in (B),(D) and (F) shown as means ± SEM compared to carrier + noise by two-way ANOVA with Bonferroni post-hoc test. *p<0.05, ***p<0.001.

Given that traumatic noise exposures are frequently unplanned, we also decided to test if oseltamivir was otoprotective when administered more than 24 hours after noise exposure. In Figure 7C & D, results are shown for mice that received 100 mg/kg oseltamivir therapy twice daily for three days beginning 48 hours after noise exposure. No protection from ABR threshold shift was observed at any frequency tested, indicating that treatment must be initiated at or within 24 hours of acoustic trauma to be protective. Finally, we assessed oseltamivir’s efficacy following an intensified noise exposure challenge of 106 dB (Figure 7E & F). Mice who received oseltamivir following exposure to 106 dB did not show reduced ABR threshold shifts relative to carrier-treated noise exposed mice.

**Figure 7:**
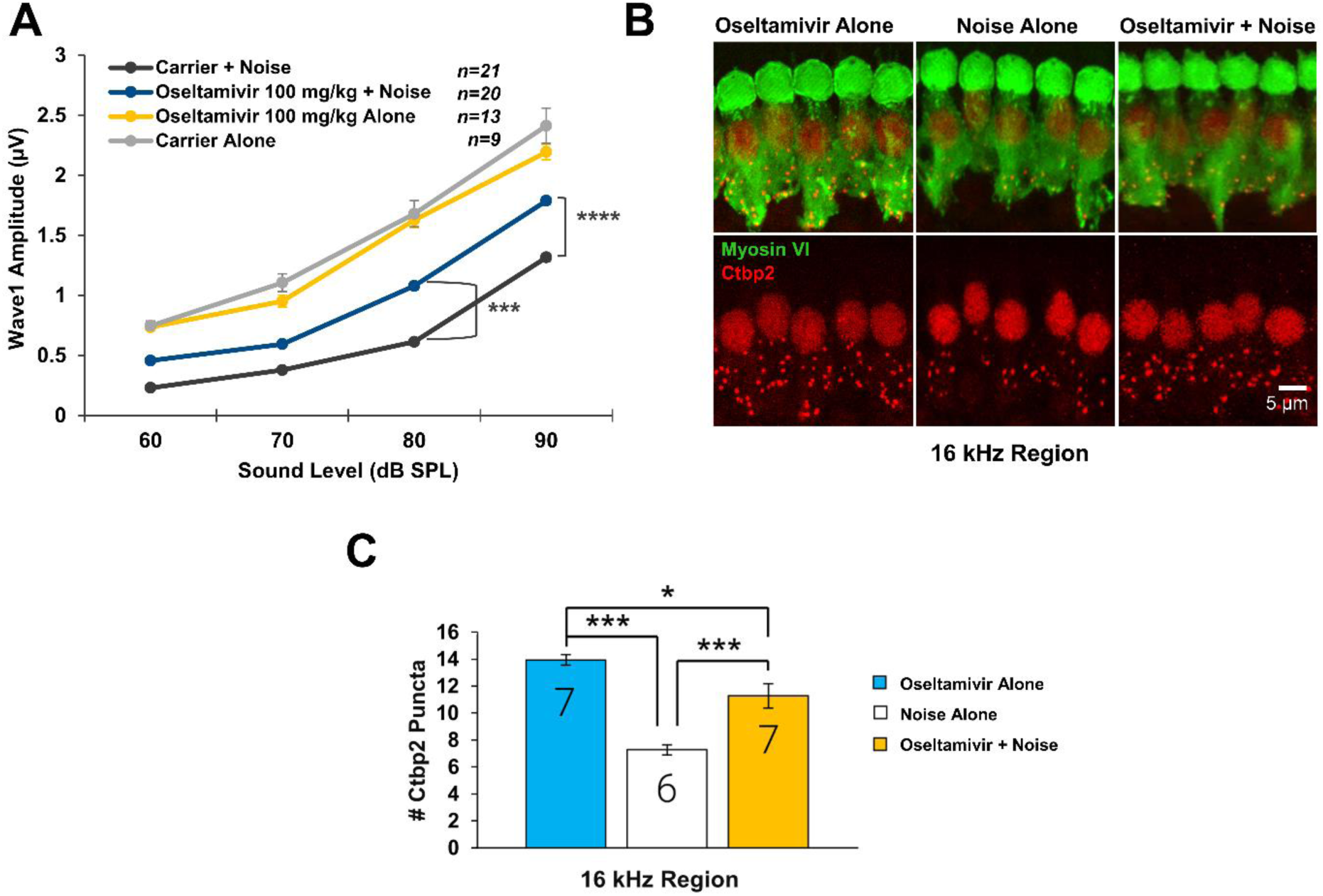
At dose of 100 mg/kg, oseltamivir protects the morphology and function of the auditory nerve synapse. **(A)** ABR wave 1 amplitude for 100 mg/kg oseltamivir vs carrier treated mice, 16 kHz. **(B, C)** Quantification and imaging of Ctbp2 (auditory synapse) puncta in treated and untreated mice. Data shown as means ± SEM compared to carrier + noise by two-way ANOVA with Bonferroni post-hoc test. *p<0.05, ***p<0.001.

### Oseltamivir therapy is protective against noise-induced cochlear synaptopathy

We analyzed synaptic function in noise-exposed mice by measuring post-treatment ABR wave 1 amplitude, a functional correlate of cochlear synaptopathy. In Figure 7A, mice treated with 100 mg/kg oseltamivir had significantly higher average ABR wave 1 amplitude at 90 and 80 dB relative to carrier-treated noise exposed mice. These results were confirmed through quantification of Ctbp2 inner hair cell synaptic puncta (Figure 7B & C). Noise-exposed mice that received 100 mg/kg oseltamivir averaged 11 Ctbp2-positive puncta per inner hair cell, significantly higher than carrier-treated noise exposed mice (7 puncta per IHC average).

### Molecular activity of oseltamivir in mitigating ototoxic effects of cisplatin is partially through inhibition of pERK protein levels and not through inhibition of mammalian neuramindases

We attempted next to determine oseltamivir’s molecular target of action in providing otoprotection from cisplatin and noise exposure. Given the known binding of oseltamivir to viral neuraminidase, we tested whether the drug would inhibit the activity of mammalian neuraminidases as well. Cochlear explants were treated with N-acetyl-2,3-dehydro-2-deoxyneuraminic acid (DANA), a pan-selective mammalian neuraminidase inhibitor, or zanamivir, an older influenza antiviral drug known to have greater off-target affinity for mammalian neuraminidases than oseltamivir (Hata et al., 2008). The drugs of interest were administered alone in medium or in combination with 150 μM cisplatin as previously described (Figure 1). When administered in combination with cisplatin, neither DANA nor zanamivir reduced OHCs death relative to cisplatin alone (Figure 8A-D). There was also no significant difference in OHCs count between explants cultured in untreated media and drug alone (DANA or zanamivir) treated explants, suggesting the low OHCs counts observed in samples cotreated with cisplatin and either DANA or zanamivir is not attributable to compound toxicity, but instead a lack of efficacy in preventing cisplatin-induced OHCs death. This result indicates that oseltamivir’s otoprotection is not mediated through neuraminidase inhibition, but through another yet undescribed target.

**Figure 8:**
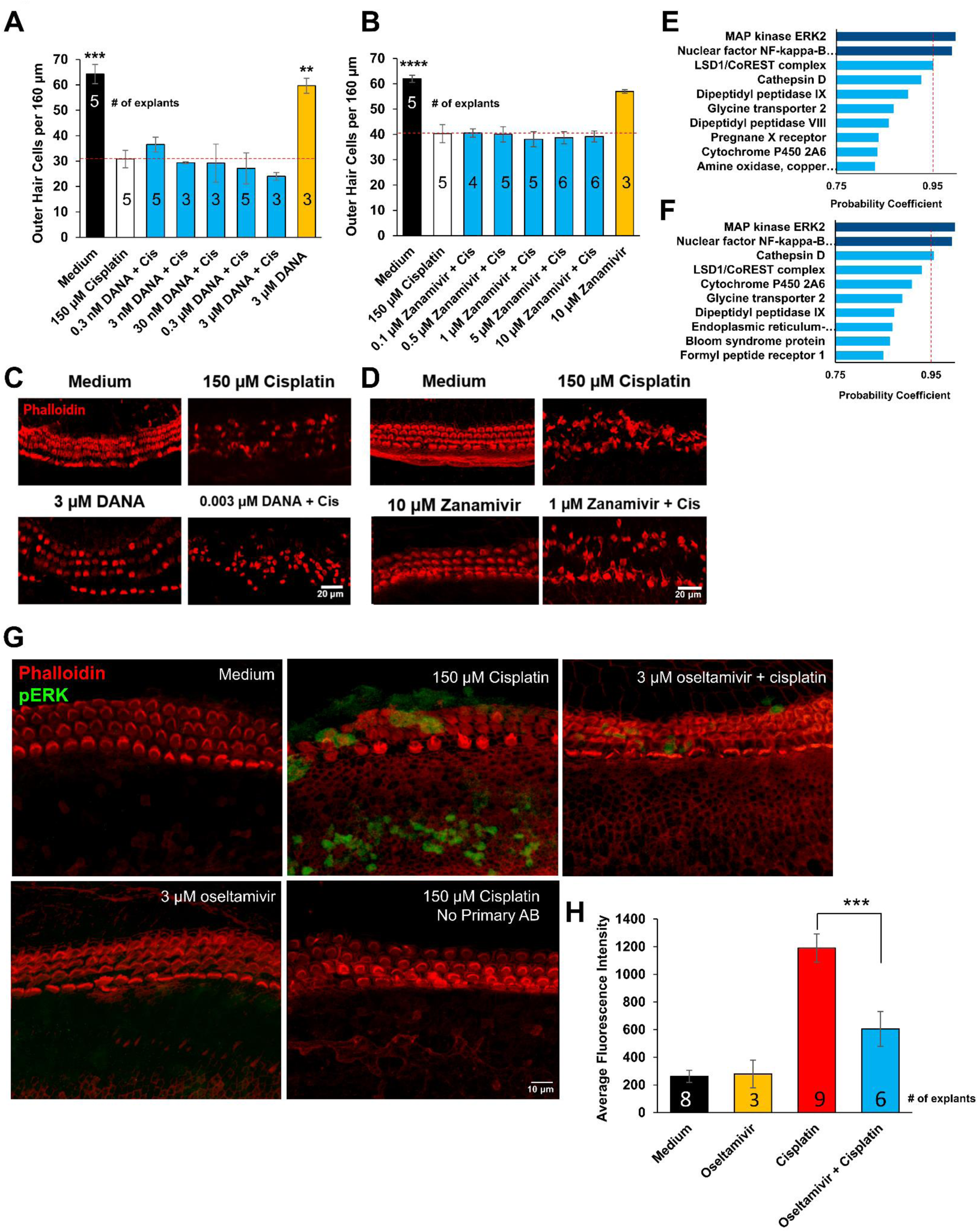
Molecular targets for oseltamivir in otoprotection and anti-inflammatory role in mice-Treatment with other neuraminidase inhibitors fails to mitigate outer hair cell loss in cisplatin-treated murine cochlear explants, indicating oseltamivir’s otoprotective qualities are mediated through non-neuraminidase targets as pERK. **(A, C)** Dose response relationship between 2,3-dehydro-2-deoxy-N-acetylneuraminic acid (DANA) and OHC count in cisplatin-treated murine cochlear explants; in (C) representative confocal images. DANA is a known inhibitor of mammalian neuraminidases NEU1-4 (IC50 = 143, 43, 61, and 74 nM, respectively). **(B, D)** Dose-response relationship between zanamivir and OHCs count in cisplatin-treated explants; in (D), representative confocal images. Zanamivir is an inhaled influenza neuraminidase inhibitor reported to weakly inhibit mammalian neuraminidases NEU1-4 (IC50 = 2,700 μM, 16.4 μM, 6.8 μM, and 487 μM, respectively). Data shown are means ± SEM, \*\**p* < 0.01, \*\*\**p* < 0.001 compared to cisplatin alone (white) by one-way analysis of variance (ANOVA) with Bonferroni post hoc test. **(E)** Top ten targets for the pro-drug, oseltamivir phosphate, predicted by drug target prediction server SuperPred. **(F)** Top ten targets predicted by SuperPred for the antiviral active metabolite, oseltamivir carboxylate. **(G)** Representative confocal images of cochlear explants immunostained with antibody anti-pERK1/2 (green) and phalloidin (red) following treatment with medium alone, 150 μM cisplatin, or 3 μM oseltamivir phosphate + 150 μM cisplatin. **(H)** Mean corrected total cell fluorescence (CTCF) of pERK1/2 expressing cells in explants following treatment with medium alone, 150 μM cisplatin in medium, or 3 μM oseltamivir phosphate in combination with 150 μM cisplatin. Data shown as means ± SEM compared to cisplatin by one-way ANOVA with Bonferroni post-hoc test. ***p<0.001.

As oseltamivir is specifically designed to target the neuraminidase activity of influenzas A and B, it does not have known molecular targets in mammals. Given the dearth of literature on its potential mammalian, nonviral target effects, we submitted the chemical structures of the pro-drug, oseltamivir phosphate, and its active metabolite, oseltamivir carboxylate, to the drug target prediction server, SuperPRED,, to generate a list of probable targets based on ATC drug class on ChEMBL binding data (Gallo et al., 2022; Nickel et al., 2014). ERK2 and NF-kB p105 were reported as top hits for both compounds at a probability coefficient exceeding 0.95 for each (Figure 8E & F).

The top two predicted targets, ERK1/2 and NF-κB, have both previously been implicated in cisplatin- and noise-induced hearing loss. ERK phosphorylation is known to be upregulated in the inner ear following both insults, as is NF-κB activation and subsequent immune cell recruitment (Ingersoll et al., 2024, 2023, 2020; Kim et al., 2008; Lutze et al., 2023; Masuda et al., 2006; So et al., 2007; Zhang et al., 2019). We therefore sought to determine if oseltamivir reduced ERK activation by studying its effects on expression of phosphorylated ERK1/2 (pERK) in cisplatin-treated P3 cochlear explants. Cochlear explants were collected and cultured as previously described and then treated with medium alone, 3 µM oseltamivir phosphate alone, 150 µM cisplatin in medium, or 3 µM oseltamivir phosphate combined with 150 µM cisplatin. Total cisplatin treatment time was 10 minutes for samples to be stained for pERK based on our previous published time-course experiments in this assay (Ingersoll et al, 2020). Mean pERK fluorescence was significantly elevated in samples receiving cisplatin alone relative to medium alone controls (Figure 8G & H), consistent with prior literature. Concurrently, treatment with 3 µM oseltamivir phosphate significantly reduced fluorescence of pERK relative to 150 µM cisplatin alone (Figure 8G & H).

### Oseltamivir reduces CD45 positive immune cells in the cochleae of noise-exposed mice

To test whether oseltamivir decreases inflammation following noise exposure, mice were exposed to 100 dB SPL noise for 2 hours with the experimental protocol listed in Figure 9A. Mouse cochleae were then collected 1 hour following the final oseltamivir treatment, approximately 4-days post noise exposure. Mouse cochleae were then sectioned and stained with CD45 (immune cell marker) and DAPI (nuclear stain). Noise exposed mice had a significantly higher average of CD45 positive cells per cochlear section (37) compared to non-noise exposed carrier (16) or oseltamivir (14) treated mice. Mice treated with 100 mg/kg oseltamivir for 3 days, twice a day had a significant reduction in the number of CD45 positive cells per cochlear section (25) compared to noise alone mice (Figure 9B & C).

**Figure 9:**
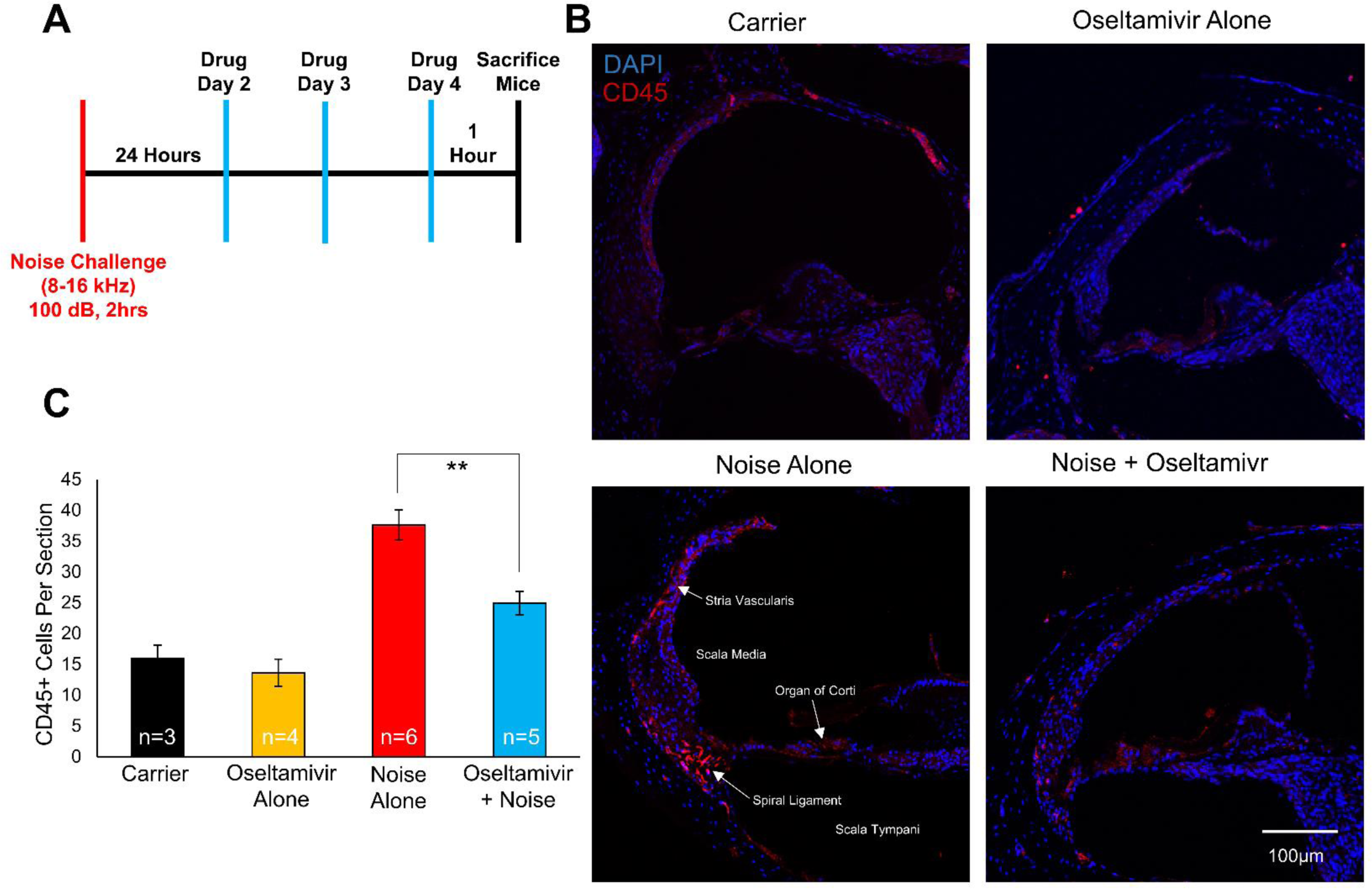
Oseltamivir decreases the number of CD45 positive cells following noise insult. **(A)** Noise exposure and treatment schedule. Mice were exposed to 100 dB noise for 2 hours (8-16 kHz) and treated with oseltamivir for 3 days starting 24 hours following noise exposure. Cochleae were collected 4 days post noise insult. **(B)** Representative confocal images of cochlear cryosections stained with CD45 (red) and DAPI (blue). **(C)** Quantification of CD45 positive cells per cochlear section. Four treatment groups are carrier alone (black), oseltamivir alone (yellow), noise + carrier (red), and oseltamivir + noise (blue). Data shown as means ± SEM, **P<0.01 compared to noise alone by one-way ANOVA with Bonferroni post hoc test. n=3-6 mice.

## Discussion

We found that oral oseltamivir therapy protected functional measures of hearing in the setting of both cisplatin and noise-induced hearing loss. This protection was demonstrated in two different mouse strains, FVB/NJ and CBA/CaJ, in both female and male mice. Oseltamivir protects from cisplatin ototoxicity in a single, high dose cisplatin treatment model employing FVB/NJ mice. Since patients receive multiple low doses of cisplatin over several treatment cycles, we then confirmed the drug’s efficacy against cisplatin ototoxicity in a more clinically relevant multi-cycle cisplatin treatment model using CBA/CaJ mice. In this multi-cycle model of cisplatin administration, both 50 and 10 mg/kg oral oseltamivir treatment significantly reduced ABR threshold shifts and protected outer hair cell count relative to cisplatin alone. Similarly, we performed noise exposure experiments in FVB/NJ mice and found significant protection from elevations in ABR threshold shifts in this strain. When given up to 24 hours after a traumatic noise exposure, 100 mg/kg oseltamivir reduced ABR threshold shifts by an average of 20-25 dB in all frequencies tested, while mice treated with 50 mg/kg showed significant average reduction of 20 dB in the 8 kHz region. Moreover, oseltamivir treatment significantly protected ABR wave 1 amplitude in cisplatin-exposed mice, as well as Ctbp2 puncta count in noise-exposed mice. This indicates that it prevents cochlear synaptopathy as a potential functional mechanism of action for hearing protection.

In the settings of both cisplatin and noise exposure, oseltamivir-treated mice exhibited significantly lower ABR threshold shifts compared to untreated mice. However, oseltamivir did not significantly reduce average DPOAE threshold shifts. DPOAE is a measurement of outer hair cell function, while ABR measures neuronal activity along the auditory pathway from the auditory nerve to the inferior colliculus located in the brainstem (Abdala and Visser-Dumont, 2001; Eggermont, 2019; Young and Ng, 2024). This indicates that the drug does not prevent changes in outer hair cell function secondary to cisplatin or noise exposure. Rather, it may protect auditory nerve function and/or central auditory processing. On a functional level, our results suggest one major mechanism of otoprotection conferred by oseltamivir is prevention of cochlear synaptopathy. Cochlear synaptopathy, the loss of synapses between inner hair cells and spiral ganglion neurons, is both a frequent consequence of traumatic noise exposure and a cause of noise-induced hearing loss (Kobel et al., 2017). In this study, we found that oseltamivir treated mice had both significantly higher ABR wave 1 amplitudes, a functional measure of synaptic health (Liberman and Kujawa, 2017), as well as significantly higher Ctbp2 puncta counts, an inner ear synaptic ribbon marker (Paquette et al., 2016), following exposure to acoustic trauma compared to noise-exposed mice that received carrier alone. These results suggest that oseltamivir treatment mitigated or prevented insult to cochlear synapses in the context of acoustic trauma, and thereby prevented increases in ABR threshold shifts that may occur secondary to synaptopathy. Oseltamivir could be protecting from synaptopathy by decreasing inflammation following noise insult, as demonstrated in Figure 9; however, future studies will investigate this further.

Given oseltamivir’s status as a selective viral neuraminidase inhibitor, with no known significant off-target effects in mammalian proteins, its molecular mechanism of action as an otoprotective compound is not obvious (He et al., 1999). Since mammalian neuraminidases, also known as sialidases, are variously involved in fine regulation of cell signaling throughout the body, we hypothesized that oseltamivir was exerting off-target inhibition on these innate neuraminidases, effecting downstream signaling changes that are otoprotective (Heimerl et al., 2022). However, we found that the panselective mammalian neuraminidase inhibitor, DANA, failed to prevent cisplatin-induced hair cell death in cochlear explants (Figure 8A and C). Zanamivir, an older antiviral neuraminidase inhibitor previously shown to have greater affinity for mammalian neuraminidases than oseltamivir (Hata et al., 2008), was also ineffective in this setting (Figure 8B and D). These results suggest that inhibition of mammalian neuraminidases is not otoprotective in the setting of cisplatin treatment, with oseltamivir acting instead on a currently unknown target.

The literature on potential nonviral targets of oseltamivir primarily concerns its binding affinity and inhibitory efficacy for the mammalian neuraminidases NEU1—4, with both binding affinity and inhibitory efficacy reported to be low (Hata et al., 2008). Given the lack of data available on oseltamivir’s potential interactions on other nonviral pharmacological targets, we ran the Super-PRED drug target prediction utility to generate a list of predicted targets based on chemical structure for both the pro-drug, oseltamivir phosphate, as well as the active form, oseltamivir carboxylate (Gallo et al., 2022; Nickel et al., 2014). Interestingly, Super- PRED reported ERK and NF-kB as top hits for both forms, at probabilities respectively ranging from 99.35— 99.65% and 98.69-98.84% (Figure 8 E-F). Our lab has previously reported that inhibitors of ERK, and upstream MAP kinases, including BRAF and MEK, ameliorate hearing loss in mouse models of noise and cisplatin exposure (Ingersoll et al., 2024, 2023, 2020; Lutze et al., 2023). Moreover, we and others have found that pERK1/2 expression is upregulated in the inner ear following either insult (Alagramam et al., 2014; Herranen et al., 2018; Ingersoll et al., 2024, 2023, 2020; Lahne and Gale, 2008; Maeda et al., 2013). Similarly, NF-kB activation within the inner ear has been reported in the setting of both cisplatin cytotoxicity and acoustic trauma, correlating with inflammation and immune cell recruitment (Kim et al., 2008; Masuda et al., 2006; So et al., 2007; Zhang et al., 2019). The results of our study show that pERK1/2 levels were reduced in oseltamivir co-treated cochlear explants with cisplatin compared to cisplatin alone treated explants, indicating that oseltamivir does reduce MAPK activation as predicted by the Super-PRED program. Additionally, the immune cell infiltration was reduced following noise exposure with oseltamivir treatment which indicates that oseltamivir also reduces inflammation following noise insults. Interestingly, several studies have shown that there is increased antiviral activity when MEK inhibitors are combined with oseltamivir compared to either one alone (Droebner et al., 2011; Haasbach et al., 2017, 2013). This suggests that oseltamivir acts on ERK1/2 as a previously unreported nonviral target, and moreover indicates it inhibits MAPK signaling within the inner ear as its otoprotective mechanism of action.

Oseltamivir reduced the number of CD45 positive immune cells in the cochleae of noise-exposed mice compared to mice exposed to noise without treatment showing the drug has an anti-inflammatory role in vivo. The biggest increases in these CD45 positive cells following noise exposure were in the spiral ligament, stria vascularis, and walls of the scala tympani. Oseltamivir treatment decreased the number of immune cells in these locations. Our laboratory and others have demonstrated that these areas of the cochlea have increases in immune cell infiltration following noise exposure (Bae et al., 2021; Hirose et al., 2005; Lutze et al., 2023; Tornabene et al., 2006). Oseltamivir treatment lowered the number of immune cells and brought it close to the number of immune cells in the control mice not exposed to noise. This suggests that oseltamivir decreases the inflammatory environment which has been shown to contribute and exacerbate hearing loss. Oseltamivir has been shown to have anti-inflammatory effects in cell lines, mice, and humans in a context other than hearing loss (Liu et al., 2024; Nomura et al., 2017; Wong et al., 2011). Other laboratories have shown that targeting the inflammatory response following noise insult or cisplatin administration can protect from hearing loss (Al Aameri et al., 2023; Dickey et al., 2005; Frye et al., 2019; Li et al., 2021; Orgel et al., 2023; Ramkumar et al., 2021). This could be part of the mechanism in which oseltamivir is protecting mice from both noise trauma and cisplatin ototoxicity.

In summary, we have shown that oseltamivir effectively reduces hearing loss in multiple mouse models of both noise exposure and cisplatin treatment. Oseltamivir is advantageously positioned for repurposing as an ototherapeutic. As a first-line influenza antiviral, it has been used in large patient populations for decades and is widely available around the world (Davies, 2010; Dutkowski et al., 2003; McClellan and Perry, 2001). Given the readiness of generic formulations, it is a relatively inexpensive drug. In our studies, mice had significant protection from hearing loss when oseltamivir was administered orally, which is the most convenient route of administration for patients and providers. At the doses administered following noise exposure or cisplatin administration, no signs of toxicity were observed, consistent with prior clinical trial data of equivalent human doses. Oseltamivir was protective when given up to 24 hours after exposure, indicating that it may be effective when given after unplanned damaging noise exposures. In the setting of the translationally relevant mouse model of cisplatin ototoxicity, we observed significant protection with a dose of 10 mg/kg given twice a day, which is 66% of the mouse equivalent of the standard adult influenza dose (Nair and Jacob, 2016).

Additionally, oseltamivir did not interfere with cisplatin’s tumor killing ability *in vitro.* This is critical to demonstrate because the treatment of the tumor is the top priority for the clinician and patient. Future studies will test whether the drug interferes with cisplatin’s tumor killing ability in tumor mouse models to confirm non-interference *in vivo* (Freyer et al., 2023). The results presented in this study demonstrate promising preclinical data that oseltamivir can be repurposed to protect from both cisplatin and noise-induced hearing loss.

## Materials and Methods

### Ethics statement

All animal procedures were approved by the Institutional Animal Care and Use Committee of Creighton University (IACUC) in accordance with policies established by the Animal Welfare Act (AWA) and Public Health Service (PHS).

### Mouse models

All noise exposure experiments and the single dose cisplatin protocol were conducted using FVB/NJ mice acquired from Jackson Laboratory (Bar Harbor, Maine, USA) and descendants bred in the Creighton University Animal Resource Facility (ARF). For the multicycle cisplatin mouse model, 8-week old CBA/CaJ mice were purchased from Jackson Laboratory and were given one week to acclimate before any experiments began. For all mouse models, anesthesia was performed using 500 mg/kg Avertin (2,2,2-tribromoethanol) delivered via intraperitoneal injection. Loss of righting reflex and pedal reflex (as assessed via toe pinch) were used as benchmarks for determining whether adequate levels of anesthesia had been achieved prior to measurement of ABR and DPOAE. Mice were randomly assigned to all treatment groups with equal numbers of female and male mice included per group.

### High-throughput screen for identifying candidate drugs protective from cisplatin-induced cell death in murine inner ear cell line

Oseltamivir phosphate was identified as potentially otoprotective via the cell-based high-throughput screen described (Ingersoll et al., 2020; Teitz et al., 2018, 2016). Briefly, cells of the murine inner ear cell HEI-OC1 were plated at a previously optimized concentration of 1600 cells per well and tested against 50 μM cisplatin for apoptotic activity via the Caspase 3/7-Glo assay. Approximately 1,300 FDA approved small molecule compounds were tested in the screen where oseltamivir phosphate was identified as a top hit. The compounds were dissolved in DMSO prior to addition to medium with final DMSO concentration in medium kept below 0.5%. Pifithrin-α was employed as a positive reference compound with an IC_50_ of 7.7 μM for inhibition of cisplatin-induced caspase-3 cleavage. Hits were defined as compounds that reduced caspase-3/7 activity by 50% or more when co-administered with 50 μM cisplatin.

### Cochlear explants

Cochleae were collected from P3 FVB/NJ mice (Ingersoll et al., 2020;Teitz et al., 2018). Immediately following collection, cochleae were plated in six-well plates (2-3 explants/well) containing 1 ml media (growth medium DMEM (12430-054,Gibco Life Technologies) combined with 1% FBS (16000-044, Gibco Life Technologies), B-27 supplement (200 μl/500 ml; 17504-44, Gibco Life Technologies), N-2 supplement (100 μl/500 ml; 17502-048, Gibco Life Technologies), and ampicillin (50 μg/ml; A5354-10ML, Sigma-Aldrich). Cochlear explants were then incubated for 24 hours at 37°C and 5% CO_2_. Following 24 hours, the explants were pretreated with fresh media with or without the compounds of interest (oseltamivir phosphate, oseltamivir carboxylate, zanamivir, or DANA) at various concentrations and then incubated at 37°C and 5% CO_2_ for an additional hour. Media was then replaced with fresh growth media with or without the compounds of interest in addition to 150 μM cisplatin (479306, Sigma-Aldrich), after which the explants were again incubated for a final 24 hours at 37°C and 5% CO_2_. Cochlear explants were then fixed in 4% paraformaldehyde in 1x PBS for a minimum of one hour. The explants were stained for F-actin with Alexa Fluor 568 Phalloidin (1:400 dilution, A12380, Thermo Fisher Scientific) or antibody anti-pERK1/2 (Thr^202^/Tyr^204^, 1:400 dilution, 9101, Cell Signaling Technology) and imaged with a confocal microscope (LSM 700, Zeiss). Outer hair cell number were counted per 160 μm in the cochlear middle turn region.

### Drug preparation and administration

Oseltamivir phosphate powder was acquired from MedChemExpress (HY-17016) and dissolved in a carrier solution of 10% DMSO, 40% PEG 300, 5% Tween-80 and 45% saline for administration by oral gavage. Doses of 100, 50, 10, or 2 mg/kg were administered to the mice at the respective times for each individual experiment. Oseltamivir carboxylate, DANA and zanamivir powder was purchased from MedChemExpress (HY-13318, HY-125798, HY-13210) and dissolved in DSMO prior to serial dilution in media for the cochlear explant experiments.

### Auditory brainstem response

ABR waveforms recordings were collected in mice anesthetized as described previously (Ingersoll et al., 2024, 2023, 2020; Lutze et al., 2023; Teitz et al., 2018). Recordings occurred in a sound booth (Industrial Acoustic Company) via three subdermal needle electrodes placed medially down the skull, below the pinna of the left ear, and at the base of the tail. Responses were fed into a low-impedance Medusa digital biological amplifier system (RA4L; TDT; 20-dB gain). At 8, 16, and 32 kHz, mice were exposed to auditory stimuli starting at 90 dB; to determine the minimum threshold dB-SPL for each frequency tested, the stimulus intensity was then reduced in 10 dB intervals until a minimum intensity of 10 dB was reached. ABR waveforms were averaged in response to 500 tone bursts and recorded signals were filtered by a band-pass filter (300 Hz to 3 kHz). ABR threshold was defined as the lowest dB tested where at least 3 of the 5 characteristic waveform peaks were still present. For all experiments, mice with baseline ABR thresholds equaling or exceeding 50 dB at any frequency were considered to have pre-existing hearing loss and thus were excluded from further experimental use. Furthermore, all ABR threshold readings were independently validated by 2 to 3 additional readers blinded to the treatment group assignments of the mice. Threshold shifts were calculated by subtracting pre-exposure thresholds from post-exposure thresholds for each frequency. ABR wave 1 amplitude were measured at 16 kHz in post-exposure recordings and were defined as the difference between the wave 1 peak and the noise floor for a given ABR trace.

### Distortion product otoacoustic emission

DPOAE recordings were collected from mice anesthetized as previously described (Ingersoll et al., 2024, 2023, 2020; Lutze et al., 2023). DPOAE recordings were collected in a sound booth (Industrial Acoustic Company) using the ER10B+ microphone system, with ear tip and speaker tubes placed in the left ear canal such that the tympanic membrane remained unobstructed. Data collection and analysis was performed on the TDT RZ6 workstation running BioSigTZ software (Tucker-Davis Technologies). DPOAE was measured at 8, 16, and 32 kHz with an f2/f1 ratio of 1.2. Tone 1 was *.909 of the center frequency and tone 2 was *1.09 of the center frequency. DPOAE data was recorded every 20.97 milliseconds and averaged 512 times at each intensity level and frequency tested. For each frequency tested, the initial stimulus intensity was 90 dB and was then decreased in increments of 10 dB until a minimum intensity of 10 dB was reached. The DPOAE threshold was defined as the lowest dB tested at a given frequency to exhibit an emission above the noise floor. DPOAE threshold shift was defined as equaling the difference of the baseline DPOAE threshold and the post-experimental DPOAE threshold for each frequency tested.

### Traumatic noise exposure model and treatment

Baseline hearing tests (ABR and DPOAE) were performed as previously described on 6-8 week old FVB/NJ mice one week in advance of noise exposure. To model noise-induced hearing loss and acoustic trauma, the mice were exposed to 100 dB SPL noise over a 8-16 octave band for two hours. The noise challenge took place within a sound-tight acrylic chamber (custom-built; Creighton University physics machining shop) which housed a top-mounted JBL speaker positioned above and facing down towards the mice. To ensure all mice received equivalent noise coverage from the speaker, the sound chamber was equipped with a metal wire enclosure consisting of ten individual compartments; these compartments were distributed in a circle, with each compartment equidistant from the other as well as from the top-mounted speaker. The sound stimulus was generated using a System RZ6 (Tucker-Davis Technologies) workstation and amplified through a 75-A power amplifier (Crown). Prior to each noise exposure challenge, sound pressure level was calibrated using an NSRT-mk3 microphone (convergence instruments) to confirm each compartment within the enclosure was within 0.5 dB of 100 dB and ensure equal noise exposure for all mice. Treatment with oseltamivir phosphate (10, 50, or 100 mg/kg) or carrier began 24 hours after the cessation of noise exposure, and was continued twice daily for 3-5 days including day of first treatment; the efficacy of treatment with 100 mg/kg oseltamivir phosphate when initiated 48 hours was also studied. Post-exposure ABR and DPOAE measurements were then collected 14 days after noise exposure, when mice were 8-10 weeks of age.

### Single high dose cisplatin treatment model

Pre-experimental ABR were performed on 6-8 week old FVB mice. One week after ABR testing, mice were treated with 50 mg/kg oseltamivir phosphate 45 minutes before a 30 mg/kg cisplatin intraperitoneal injection in the morning. Mice were then treated with oseltamivir phosphate again in the evening. Oseltamivir phosphate was administered twice a day for three total days. 21 days after the single cisplatin injection, post-experimental ABR were performed, and cochleae were harvested and put in 4% PFA solution. One day before cisplatin injection, mice received 1 mL of saline by subcutaneous injection and were given 1 mL of saline twice a day throughout the protocol until body weight started to recover. The cages of cisplatin treated mice were placed on heating pads until body weights began to recover. Food pellets dipped in DietGel Boost® were placed on the cage floor of cisplatin-treated mice. DietGel Boost® (72-04-5022 Clear H2O) is a high calorie dietary supplement that provides extra calorie support for mice. The investigators and veterinary staff carefully monitored for changes in overall health and activity that may have resulted from cisplatin treatment.

### Multi-cycle cisplatin treatment model

Pre-experimental ABR were performed on 9-week old CBA/CaJ mice with DPAOE performed when mice were 10 weeks old. Once mice were 12 weeks old, the 6-week cisplatin and oseltamivir phosphate treatment regimen began. Oseltamivir phosphate (50, 10, or 2 mg/kg/bw) was administered via oral gavage 1 hour before 3 mg/kg cisplatin was administered to mice via intraperitoneal injection in the morning. Mice were then treated with oseltamivir phosphate or carrier again in the evening. Mice were treated with cisplatin once a day for 4 days and oseltamivir phosphate twice a day for 5 days with a 9-day recovery period in which no drugs were administered to the mice. This cycle was repeated two more times for a total of 3 cycles. Mice were treated with 3 mg/kg cisplatin for a total of 12 days (4 days per cycle with 3 cycles) and oseltamivir phosphate for a total of 15 days (5 days per cycle with 3 cycles). Immediately after the completion of cycle 3 (42 days after the first cisplatin injection), post-experimental ABR were performed with DPOAE performed one week after ABR. Cochleae were when harvested and put in 4% PFA. Cisplatin treated mice were injected by subcutaneous injection twice a day with 1 mL of warm saline to ameliorate dehydration. This continued until body weight started to recover. The cages of cisplatin-treated mice were placed on heating pads throughout the duration of the experiment until mice began to recover after the 3rd treatment cycle of the protocol. Food pellets dipped in DietGel Boost® were placed on the cage floor of cisplatin-treated mice. The investigators and veterinary staff carefully monitored for changes in overall health and activity that may have resulted from cisplatin treatment.

### Outer hair cell counts

Cochleae from adult mice were prepared and examined as described previously (Ingersoll et al., 2023; Yamashita et al., 2012). Cochleae samples were immunostained with anti-myosin VI (1:400; 25-6791, Proteus Bioscience) with secondary antibodies purchased from Invitrogen coupled to anti-rabbit Alexa Fluor 488 (1:400; A11034). All images were acquired with a confocal microscope (LSM 700 or 710, Zeiss). Outer hair cell counts were determined by the total amount of OHCs in a 160 µm region. Counts were determined for the 8, 16, and 32 kHz regions. Cochleae from each experimental group were randomly selected to be imaged for outer hair cell counts.

### Ctbp2 staining for quantification of auditory nerve puncta

Cochlear dissections were performed at the conclusion of noise exposure studies. After fixation in 4% PFA, otic capsules underwent brief decalcification for 30 minutes in a 120 mM EDTA solution. The organs of corti were then dissected from the otic capsule and co-stained with anti-Ctbp2 antibodies (1:800; 612044, BD Transduction) and anti-myosin VI antibodies (1:400; 25-6791, Proteus Biosciences) overnight at 4°C. Goat anti-rabbit Alexa Fluor 488 (1:400; A11034) and goat anti-mouse Alexa Fluor 647 (1:800; A32728) were purchased from Invitrogen as the secondary antibodies. Images of the 16 kHz regions of each sample were collected upon the LSM 700 confocal microscope (Zeiss) at a 63x objective. Maximum intensity projections were obtained using ZEN BLACK software (Zeiss) with Ctbp2 puncta counted in ImageJ. 12-18 inner hair cells were visible per image and average number of Ctbp2 puncta per sample was calculated as equaling the total number of puncta divided by the total number of inner hair cells visible. Cochleae used for Ctbp2 puncta quantification were obtained from 6-7 randomly selected mice per treatment group.

### Cochlear cryosectioning and CD45 staining

FVB mice aged 6-8 weeks’ old were exposed to 100 dB SPL noise (8-16 kHz octave band) for 2 hours. Mice were either treated with carrier alone or 100 mg/kg oseltamivir twice a day for 3 days beginning 24 hours after noise exposure. Mice were then sacrificed 1 hour after the last drug treatment which was approximately 4 days following noise exposure. Cochleae were extracted from mice and placed in 4% PFA for 2-3 days. Cochleae were then decalcified in 120 nM EDTA for 2-3 days. Following decalcification, cochleae were transferred to a 30% sucrose solution and kept at 4°C overnight. The next day, samples were put into a solution of 30% sucrose and OCT compound (4583; Sakura) for 4 hours at 4°C. Samples were then placed in OCT compound overnight at 4°C. The next day, cochlear tissues were oriented in within cryomolds containing OCT compound and frozen on dry ice. Frozen tissues were cut with 10µm thickness. captured on glass slides and allowed to dry for several hours.

Cochlear cryosections were then blocked and permeabilized in a solution of 5% FBS and 0.2% triton X-100 in PBS. Tissues were stained overnight at 4°C with mouse CD45 antibody (1:50; Af114, R&D Systems). The next day, tissues were stained for one and a half hours with Alexa Fluor 568 donkey anti-goat (1:400; A11057, Invitrogen) and DAPI (1:1000; D1306, Invitrogen) to counterstain nuclei. Tissues were mounted in Fluoromount-G (00-4958-02, Invitrogen) and imaged using a Zeiss 700 upright confocal microscope. Post-acquisition images were analyzed using the IMARIS imaging software and automatically quantified following intensity thresholding. CD45 positive cells were then cross-checked manually to ensure positive CD45 cells were co-stained with DAPI.

### Statistical analysis

Statistical analysis was performed in Prism (GraphPad Software). Two-way analysis of variance (ANOVA) or one-way ANOVA with Bonferroni post hoc test was used to determine mean difference and statistical significance.

## Acknowledgements

We thank Daniel Kresock, Dr. Christy Howe, Dr. Janee Gelineau-van Waes, Dr. Kristen Drescher, Pat Steele, Ann Bryen, and the Creighton University ARF staff for assistance with the mouse studies. The research was funded by the following grants: Department of Defense Award # W81XWH-21-1-0696, grant RH200032, LB 506 Award from Nebraska State, Department of Health and Human Services, Cancer and Smoking Disease Research Program, National Institutes of Health NIDCD grant 1R01DC018850, American Hearing Research Foundation 2020 grant to Tal Teitz. This investigation was conducted in facilities constructed with support from Research Facilities Improvement Program (G20 RR024001-01) from the National Center for Research Resources, NIH. The research was partially conducted at the Auditory and Vestibular Technology Core (AVT) at Creighton University, Omaha, NE (RRID:SCR_023866). This facility is supported by the Creighton University School of Medicine and grants GM103427 and GM139762 from the National Institute of General Medical Science (NIGMS), a component of the National Institutes of Health (NIH). IBIF was constructed with support from grants from the National Center for Research Resources (RR016469) and the NIGMS (GM103427). This investigation is solely the responsibility of the authors and does not necessarily represent the official views of the National Center for Research resources, NIGMS or NIH.

## Author Contributions

T.T., D.C., J.Z. and T.C. designed and performed the cell-based screens, M.A.I. and R.D.L. performed the in vivo cisplatin experiments. E.L. and R.G.K., performed the in vivo noise exposure experiments. M.A.I. performed the cochlear dissections, outer hair cell, and Ctbp2 counts. E.L., M.A.I., and K.L. performed cochlear explants experiments, E.L., R.G.K., M.A.I., and R.D.L., analyzed ABR and DPOAE data, R.D.L. and R.G.K. performed and imaged the cochlear CD45 stained cryosections. R.D.L. performed the cisplatin interference testing in tumor cell lines. E.L., M.A.I., R.D.L., R.G.K., K.L., and T.T. were involved in analysis of data and the design of the study. E.L., R.D.L., K.L., R.G.K., M.A.I., and T.T. wrote the manuscript with input from all authors.

## Data, Materials, and Software Availability

All data needed to evaluate the conclusions in the paper are present in the paper and/or Supplementary Materials.

